## Supplementary Information for "Influence of On-Off dynamics and selective attention on the spatial pattern of correlated variability in neocortex"

**4 Supplementary figures 30**

### 1 Data analysis

#### 1.1 Multi- and single-unit activity

Multi- and single-unit activity (MU and SU) detection procedures have been described in previous study [22]: The raw 40 kHz-sampled voltage trace was match-filtered and peaks were detected. Then, a moving-threshold algorithm was employed with a very high threshold to detect the largest peaks and to exclude any small spikes that occurred within the exclusion window around these largest peaks. For MUs, progressively lower thresholds were used until approximately 100 Hz of spiking activity was detected on each channel. For SUs, out of 736 total recorded channels in the 46 sessions, we were able to isolate 285 single neurons. We use MU activity for HMM-fitting. Attentional modulation of firing rates, Fano factor, and noise correlations was similar for MU and SU activity (Supplementary Fig. 2, Supplementary Tables 1,2).

#### 1.2 Attentional modulation of firing rate, Fano factor and noise correlations in one-phase and two-phase recordings

We quantified attentional modulation of firing rates, Fano factor (FF) and noise correlations in one-phase and two-phase recordings, separately for MU and SU in superficial and deep cortical

| | | Layer | MI | $p$-value | $n$ |
|---|---|---|---|---|---|
| Firing rate | MU | Superficial | 0.0225 | $p < 10^{-10}$ | $1,752$ |
| | MU | Deep | 0.0183 | $p < 10^{-10}$ | $2,216$ |
| | SU | Superficial | 0.0217 | $p < 10^{-9}$ | 944 |
| | SU | Deep | 0.0167 | $p < 10^{-6}$ | $1,282$ |
| Fano factor | MU | Superficial | $-0.0101$ | $p < 10^{-10}$ | $1,752$ |
| | MU | Deep | $-0.0067$ | $p < 10^{-10}$ | $2,216$ |
| | SU | Superficial | $-0.0013$ | $p = 0.30$ | 944 |
| | SU | Deep | $-0.0014$ | $p = 0.26$ | $1,282$ |
| Noise correlation | MU | Superficial | $-0.0294$ | $p < 10^{-5}$ | $5,088$ |
| | MU | Deep | 0.0215 | $p = 0.004$ | $6,128$ |
| | SU | Superficial | $-0.0431$ | $p = 0.1$ | $2,011$ |
| | SU | Deep | 0.0768 | $p < 0.001$ | $4,112$ |

**Table 1. Attentional modulation of firing rate, Fano factor, and noise correlations in two-phase recordings.** Modulation index (MI) is computed as the difference between attention and control conditions divided by the sum. $p$-value is for Wilcoxon rank-sum test. For firing rate and Fano factor, $n$ is the number of units. For noise correlation, $n$ is the number of unit pairs.

| | | Layer | MI | $p$-value | $n$ |
|---|---|---|---|---|---|
| Firing rate | MU | Superficial | 0.0241 | $p < 10^{-10}$ | 904 |
| | MU | Deep | 0.0307 | $p < 10^{-10}$ | $1,016$ |
| | SU | Superficial | 0.0204 | $p = 0.005$ | 324 |
| | SU | Deep | 0.0448 | $p < 10^{-10}$ | 532 |
| Fano factor | MU | Superficial | $-0.0077$ | $p < 10^{-10}$ | 904 |
| | MU | Deep | $-0.0098$ | $p < 10^{-10}$ | $1,016$ |
| | SU | Superficial | 0.0001 | $p = 0.75$ | 324 |
| | SU | Deep | $-0.0069$ | $p = 0.0045$ | 532 |
| Noise correlation | MU | Superficial | $-0.0306$ | $p = 0.10$ | 920 |
| | MU | Deep | 0.0628 | $p = 0.0003$ | $1,448$ |
| | SU | Superficial | $-0.0208$ | $p = 0.53$ | 421 |
| | SU | Deep | $-0.018$ | $p = 0.41$ | 901 |

**Table 2. Attentional modulation of firing rate, Fano factor, and noise correlations in one-phase recordings.** Same format as in Supplementary Table 1.

layers (Supplementary Tables 1,2). We calculated a standard modulation index $MI_{rate}$ ($MI_{FF}$, $MI_{corr}$), which was the difference between the firing rate (FF, noise correlation, respectively) in the attention and control conditions divided by the sum.

### 2 Network model of interacting columns

#### 2.1 On-Off dynamics in single columns

On-Off dynamics of individual units in the model describe synchronized transitions in population neural activity within cortical columns. A group of neurons within a column (indexed by $i = 1, 2...$) simultaneously switches between On and Off phases, which is characterized a binary value $S(t) = \{0, 1\}$, where $S(t) = 1$ denotes On phase and $S(t) = 0$ denotes Off phase. The neurons have higher firing rates $r_{\text{on}}(i)$ during On phases, and lower firing rates $r_{\text{off}}(i)$ during Off phases, which can be represented by the time-dependent firing-rate $\lambda(i; t)$:

$$\lambda(i;t) = r_{\text{off}}(i) + S(t)[r_{\text{on}}(i) - r_{\text{off}}(i)] \ . \tag{1}$$

$S(t)$ transitions between On and Off phases stochastically as a two-state Markov process. The transition rate from Off to On phase is $\alpha_1$, and the transition rate from On to Off phase is $\alpha_2$. Therefore, the mean duration of Off episode is $\tau_{\text{off}} = 1/\alpha_1$, and the mean duration of On episode is $\tau_{\text{on}} = 1/\alpha_2$. The spike counts of neurons are generated as inhomogeneous Poisson processes with instantaneous firing rates $\lambda(i; t)$.

##### 2.1.1 Analytical prediction of Fano factor and noise correlations in single columns

In the model, columnar On-Off dynamics are the source of correlated variability within single model units. Spike counts of each neuron within a column are generated by a doubly stochastic process, where shared On-Off dynamics give rise to noise correlations between neurons. We tested how accurately the model of On-Off dynamics predicted attentional modulation of FF and noise correlations in our columnar recordings. We analytically calculated FF and noise correlation within an arbitrary time window $T$ as a function of the On-Off transition rates ($\alpha_1$ and $\alpha_2$) and On and Off firing rates ($r_{\text{on}}$ and $r_{\text{off}}$) of individual neurons.

On each trial, the spike-count $N(i)$ of a neuron $i$ in the time-window $T$ is described by a Poisson random variable with the rate given by the integral

$$\Lambda(i) = \int_t^{t+T} \lambda(i; t')dt' = r_{\text{off}}(i)T + R\Delta r(i) \ . \tag{2}$$

Here the On-Off firing-rate difference $\Delta r(i)$ is defined as

$$\Delta r(i) = r_{\text{on}}(i) - r_{\text{off}}(i) \ , \tag{3}$$

and $R$ is the normalized rate:

$$R = \int_{t}^{t+T} S(t')dt' \ . \tag{4}$$

FF is defined as the ratio of the spike-count variance to the mean:

$$FF_i = \frac{\text{Var}[N(i)]}{\text{E}[N(i)]} \ . \tag{5}$$

Noise correlation between spike counts of neurons $i$ and $j$ is defined as

$$NC_{i,j} = \frac{\text{Cov}[N(i), N(j)]}{\sqrt{\text{Var}[N(i)]\text{Var}[N(j)]}} = \mathcal{A}(\alpha_1, \alpha_2) \ , \tag{6}$$

where $\text{Cov}[N(i), N(j)]$ is the covariance between spike-counts in a pair of neurons.

The mean value of the spike-count of neuron $i$ is given by

$$\text{E}[N(i)] = \langle \sum_{n=1}^{\infty} n \frac{(\Lambda(i))^n}{n!} e^{-\Lambda(i)} \rangle = \langle \Lambda(i) \rangle = r_{\text{off}}(i)T + \text{E}[R]\Delta r(i) \ . \tag{7}$$

The variance of spike count is

$$\begin{aligned} \text{Var}[N(i)] &= \text{E}[N(i)^2] - (\text{E}[N(i)])^2 = \langle \sum_{n=1}^{\infty} n^2 \frac{(\Lambda(i))^n}{n!} e^{-\Lambda(i)} \rangle - (\langle \Lambda(i) \rangle)^2 \\ &= \langle [\Lambda(i)]^2 + \Lambda(i) \rangle - (\langle \Lambda(i) \rangle)^2 \\ &= (\Delta r(i))^2 \text{Var}[R] + r_{\text{off}}(i)T + \text{E}[R]\Delta r(i) \ . \end{aligned} \tag{8}$$

The spike-count covariance $\text{Cov}[N(i), N(j)]$ is calculated as

$$\begin{aligned} \text{Cov}[N(i), N(j)] &= \text{E}[N(i)N(j)] - \text{E}[N(i)]\text{E}[N(j)] \\ &= \langle \sum_{n=1}^{\infty} n \frac{(\Lambda(i))^n}{n!} e^{-\Lambda(i)} \sum_{m=1}^{\infty} m \frac{(\Lambda(j))^m}{m!} e^{-\Lambda(j)} \rangle - \langle \Lambda(i) \rangle \langle \Lambda(j) \rangle \\ &= \langle \Lambda(i)\Lambda(j) \rangle - \langle \Lambda(i) \rangle \langle \Lambda(j) \rangle \\ &= \Delta r(\mathbf{x}, i) \Delta r(\mathbf{y}, j) \text{Var}[R] \ . \end{aligned} \tag{9}$$

According to the statistics of a two-state Markov process, the mean and variance of the normalized rate $R$ are

$$\text{E}[R(\mathbf{x})] = \frac{\alpha_1}{\alpha_1 + \alpha_2} T, \tag{10}$$

$$\text{Var}[R(\mathbf{x})] = \frac{2\alpha_1\alpha_2}{(\alpha_1 + \alpha_2)^3} \left[ T - \frac{1}{\alpha_1 + \alpha_2} \left(1 - \exp\left(-(\alpha_1 + \alpha_2)T\right)\right) \right] \ . \tag{11}$$

In terms of the mean On and Off episode durations $\tau_{\text{off}} = 1/\alpha_1$, $\tau_{\text{on}} = 1/\alpha_2$, we get

$$\text{E}[R(\mathbf{x})] = \frac{\tau_{\text{on}}}{\tau_{\text{on}} + \tau_{\text{off}}} T \ , \tag{12}$$

$$\text{Var}[R(\mathbf{x})] = \frac{2\tau_{\text{on}}^2 \tau_{\text{off}}^2}{(\tau_{\text{on}} + \tau_{\text{off}})^3} \left[ T - \frac{\tau_{\text{on}}\tau_{\text{off}}}{\tau_{\text{on}} + \tau_{\text{off}}} \left( 1 - \exp\left( -\frac{\tau_{\text{on}} + \tau_{\text{off}}}{\tau_{\text{on}}\tau_{\text{off}}} T \right) \right) \right] \ . \tag{13}$$

#### 2.2 Dynamical-system model of spatiotemporal On-Off dynamics

To quantify spatiotemporal patterns of On-Off dynamics, we constructed a network model of interacting columns. The model describes the propagation of On-Off dynamics across cortical surface within each layer (superficial or deep). Each layer in the model consists of a two-dimensional lattice of units. Each unit is represented by a dynamical variable $v(\mathbf{x}, t)$, which represents the mean firing rate of a population of neurons within one layer of a single column. The two-dimensional lateral coordinates are denoted as $\mathbf{x}$. The dynamical equations are

$$\begin{aligned} \epsilon \frac{d}{dt} v &= F(v) - u + W\nabla^2 v + I \ , \\ \frac{d}{dt} u &= gv - u + f + \sqrt{2Q}\xi \ . \end{aligned} \tag{14}$$

Here $u$ is the adaption variable, and $\xi$ is Gaussian white noise of unity intensity (here we omit the spatial index of variables $u$ and $v$). The piece-wise linear function $F(v)$ is given by

$$F(v) = \begin{cases} -1 - v \quad , & v \leq -1/2 \\ v \quad , & -1/2 < v < 1/2 \\ 1 - v \quad , & v \geq 1/2 \end{cases} \ . \tag{15}$$

The term $W\nabla^2 v$ represents interactions among nearby units, where $\nabla^2 v = \partial_x^2 v + \partial_y^2 v + 2\partial_x \partial_y v$, and $W$ is the interaction strength. The parameters are adaptation gain $g$, adaptation baseline $f$, and noise intensity $Q$. The external current $I = I_{\text{stim}} + I_{\text{attn}}$ represents the bottom-up inputs from visual stimuli and top-down attentional that act on a local group of units. $\epsilon \ll 1$ is a constant that separates the timescales of the dynamical variable $v$ and slow adaption variable $u$. We chose the parameters such that the system operated in a bistable regime [52], where the population rate $v$ stochastically switches between two stable fixed points, corresponding to the On and Off phases. Therefore, we use a binary variable $\lambda(\mathbf{x}, t)$ which segments the activity of $v(\mathbf{x}, t)$ into the On and Off phase. The firing rates of neurons within a certain column of population $\mathbf{x}$ is given by

$$r(t) = r_{\text{off}}(\mathbf{x}) + (r_{\text{on}}(\mathbf{x}) - r_{\text{off}}(\mathbf{x}))\lambda(\mathbf{x}, t) \ , \tag{16}$$

where $r_{\text{on}}(\mathbf{x})$ and $r_{\text{off}}(\mathbf{x})$ are firing rates of neurons during On and Off phases, respectively.

##### 2.2.1 Simulations of the dynamical-system model

We simulated the dynamical-system model of spatiotemporal On-Off dynamics Eq. (14), and then generated spikes from the simulated rate variable $v(\mathbf{x}, t)$. We performed simulations on a square $256 \times 256$ grid using a discretized operator $\nabla^2 v$ and the explicit Euler scheme for time-integration with a step $dt = 0.1$. We fixed the parameters $f = 0.15$, $g = 0.55$, $\epsilon = 0.05$, $W = 0.015$, $Q = 0.0025$. The noise intensity $Q$ was chosen relatively low so that the On and Off episode durations have close to exponential distributions. The external input current is $I = I_{\text{att}} + I_{\text{stim}} + f$. We chose the parameter $f = 0.15$ so that the On and Off episode durations were similar ($\tau_{\text{off}} \approx \tau_{\text{on}}$) in the control condition ($I_{\text{att}} = 0$). To study the dependence of noise correlations on the input current, we chose $I_{\text{att}}, I_{\text{stim}} = 0, 0.0016, 0.0032, 0.0048, 0.0064, 0.008$. For the examples in Fig. 5d and Fig. 6d, $I_{\text{att}} + I_{\text{stim}} = 0.0016$.

We generated spike-counts of individual simulated neurons from the population rates $v(\mathbf{x}, t)$. We sampled firing rates $r_{\text{off}}$ and $\Delta r = r_{\text{on}} - r_{\text{off}}$ of each neuron based on the distributions of $r_{\text{off}}$ and $r_{\text{on}}$ of HMM parameters obtained by fitting two-phase recordings. We fitted a two-state HMM to MU activities on 16 channels in each of 31 two-phase recordings, separately for 8 stimulus orientations and 4 attention conditions (2 attention and 2 control conditions). The HMM fits produced a distribution of $r_{\text{off}}$ and $r_{\text{on}}$ with $31 \times 8 \times 16$ pairs of firing rates in attention versus control condition (we averaged firing rates of 2 attention and 2 control conditions).

We calibrated time units in the model to match the timescales of On-Off dynamics in the data. In the calculation of noise correlations, we used the time-window $T = 0.2$ s in the data, and the number of time-steps of simulated firing-rate sequence $n_1 = 500$. This calibration is based on the fact that in the control condition ($I = f$) $\tau_{\text{off}} = \tau_{\text{on}} \approx 250$ time-steps, which we matched to the average On and Off episode durations in control condition that are about 100 ms. To obtain the Poisson rate, we transformed the rate to a binary variable: $\lambda(t) = \Theta[v(t)]$, where $\Theta$ is the Heaviside step function. The Poisson rate of each neuron is then calculated as $r(t)\Delta t = [r_{\text{off}} + \lambda(t)\Delta r]\Delta t$, where $\Delta t = T/n_1 = 0.4$ ms according to our time calibration. For each combination of $I_{\text{att}}$, $I_{\text{stim}}$, we simulated 100 trials, each with the duration of 5,000 time-steps, which in our calibration is equivalent to 2 s.

##### 2.2.2 Dependence of noise correlation within and across columns on the external input

In simulations, we investigated changes in noise correlations within and across columns for different values of $I_{\text{att}}$ and $I_{\text{stim}}$ (Fig. 5b). To calculate noise correlations within single columns, we used 121 units from a local group ($11 \times 11$ square on the grid) that received attentional input, and 121 units from an equivalent local group without attentional input (i.e. control condition, as in Fig. 2d). We simulated 100 trials to generate the sequences $\lambda(t)$ of each unit for attention and control conditions. For each unit, we then generated spike-counts of 10 simulated neurons with the same shared On-Off sequence, using the corresponding firing rates sampled from the distributions estimated by HMM. We calculated noise correlations between these simulated neurons and averaged them over all $121 \times 9 \times 10/2$ pairs.

To calculate noise correlations across columns, we generated spike-counts for one simulated neuron per unit, and used all $121 \times 120/2$ neuron pairs. We averaged noise correlations over pairs for units with lateral distance $d \leqslant 3$. In analytical approximation, this averaging is equivalent to

$$r_{\text{sc}}^{\text{control}} = \mathcal{A}_{\text{ctl}} \int_0^3 dx p(x) \exp(-x/L_{\text{ctl}}) \ , \quad r_{\text{sc}}^{\text{attention}} = \mathcal{A}_{\text{att}} \int_0^3 dx p(x) \exp(-x/L_{\text{att}}) \ , \tag{17}$$

where $p(x)$ is the probability density of units at distance $x$.

##### 2.2.3 Estimation of the effective interaction strength between columns

We examined how the effective strength of recurrent interactions between columns depends on the attentional input $I_{\text{att}}$ (Fig. 5e). Analytical calculations using the reduced binary-unit model predict that noise correlations decrease with the lateral distance $d$ exponentially $r_{\text{sc}} = \mathcal{A}\exp(-d/L)$, and the correlation length $L$ depends on the coupling strength $\beta$ as $L = [\beta/(\alpha_1 + \alpha_2)]^{1/2}$. We used these analytical equations to estimate $\alpha_1$, $\alpha_2$ and $\beta$ from simulations of the dynamical-system network. For each $I_{\text{att}}$, we estimated $\alpha_1$ and $\alpha_2$ as inverse of the mean Off and On episode durations, respectively. We estimated $L$ from simulations by fitting the dependence of noise correlations on the lateral distance with the exponential function. The effective coupling strength was then calculated as $\beta = L^2(\alpha_1 + \alpha_2)$.

##### 2.2.4 Analytical approximation of the dynamical system by a binary switching process

In the bistable regime and for low noise intensity, the dynamical-system units in the model Eq. (14) can be analytically reduced to binary switching units. The derivation is based on the timescale separation between the fast rate variable $v$ and the slow adaptation variable $u$ [52]. In the limit $\epsilon \to 0$, the equation for the rate variable $v$ in Eqs. (14) is reduced to a static nullcline equation

$$F(v) - u + h = 0 \ , \tag{18}$$

where $h = W\nabla^2 v + I$. This piece-wise linear nullcline has an inverted N shape, with the Off and On fixed points located on its left and right branches, respectively. Due to the timescale separation, the Off and On phases can be reduced to dynamics along the left and right branches, respectively, with instantaneous transitions between the branches. On the left branch, $v$ has lower activity and it corresponds to the Off phase $S_i = 0$. On the right branch, $v$ has higher activity and it corresponds to the On phase $S_i = 1$. To implement the transitions between the branches, we introduce two absorbing boundaries at the left ($v_- = -1/2$) and right ($v_+ = 1/2$) branches of the nullcline, respectively. The corresponding values of $u$ are

$$u_+ = \frac{1}{2} + h \ , u_- = -\frac{1}{2} + h \ . \tag{19}$$

Upon reaching an absorbing boundary, the dynamics are reset to the opposite branch at the initial reset point.

Substituting the static equation Eq. (18) into the equation for the adaptation variable $u$ in Eqs. (14), we get two equations for the dynamics along each of the two branches:

$$\dot{u} = g(\mp 1 - u + h) - u + f + \sqrt{2Q}\xi \; . \tag{20}$$

Here $\mp$ corresponds to the equations for the left and right branch, respectively. This system Eq. (20) is equivalent to two one-dimensional Langevin equations with two new variables [52] :

$$x = u - \frac{g(h-1)+f}{1+g} \; , \quad y = -u + \frac{g(h+1)+f}{1+g} \; . \tag{21}$$

The variables $x$ and $y$ describe the dynamics on the left and right branch, respectively. Both $x$ and $y$ are equal to zero at the On and Off fixed points of the deterministic equations Eqs. (14). Further, we define a rescaled time and noise intensity:

$$\tilde{t} = (1+g)t \; , \quad D = Q/(1+g) \; . \tag{22}$$

With this notation, equations Eq. (20) transform into two Langevin equations:

$$\begin{aligned} \dot{x} &= -x + \sqrt{2D}\xi \; , \\ \dot{y} &= -y + \sqrt{2D}\xi \; . \end{aligned} \tag{23}$$

The absorbing boundaries ($x_-$ and $y_-$) and reset points ($x_+$ and $y_+$) for these two one-dimensional Langevin equations are given by

$$\begin{aligned} x_- &= -\frac{1}{2} + h - \frac{g(h-1)+f}{1+g} \; , \quad x_+ = \frac{1}{2} + h - \frac{g(h-1)+f}{1+g} \; , \\ y_- &= -\frac{1}{2} - h + \frac{g(h+1)+f}{1+g} \; , \quad y_+ = \frac{1}{2} - h + \frac{g(h+1)+f}{1+g} \; . \end{aligned} \tag{24}$$

This reduced system operates as a two-state switching process, where the left and right branches correspond to the Off and On phases, respectively. The transition rates between the branches are defined by the inverse mean first passage times. The mean first passage time from the left branch to the right branch defines the Off-to-On transition rate $\alpha_1 = 1/\tau_{\text{off}}$ [52]:

$$\tau_{\text{off}} = \frac{1}{(1+g)}\langle T_l \rangle = \frac{1}{(1+g)r_0}\int_{x_-}^{\infty} P_x^0(x) = \frac{\sqrt{\pi}}{1+g}\int_{x_-/\sqrt{2D}}^{x_+/\sqrt{2D}} dz e^{z^2}\text{erfc}(z) \; , \tag{25}$$

The mean first passage time from the right branch to the left branch defines the On-to-Off transition rate $\alpha_2 = 1/\tau_{\text{on}}$ [52]:

$$\tau_{\text{on}} = \frac{1}{(1+g)}\langle T_r \rangle = \frac{1}{(1+g)r_0}\int_{y_-}^{\infty} P_y^0(y) = \frac{\sqrt{\pi}}{1+g}\int_{y_-/\sqrt{2D}}^{y_+/\sqrt{2D}} dz e^{z^2}\text{erfc}(z) \; . \tag{26}$$

The mean first passage times $\tau_{\text{on}}$ and $\tau_{\text{off}}$ present the average durations of the Off and On episodes, respectively.

To find an approximate analytical expression for the mean first passage time, we use the fact that the following integral can be approximated by

$$\int_{(-1/2+x)/\sqrt{2D}}^{(1/2+x)/\sqrt{2D}} dz e^{z^2} \text{erfc}(z) \approx a + b \exp(-cx) \ , \tag{27}$$

where $a$, $b$ and $c$ are functions of $D$. We consider several noise intensities $D = 4,\ 1,\ 0.4,\ 0.1,\ 0.04/1.55$, and for each $D$ perform the nonlinear fitting to find the best fit parameters:

$$\begin{aligned}
D = 4 \quad &: \quad a = 0.13203\ , \quad b = 0.22504\ , \quad c = 0.52392\ , \quad R^2 = 0.99998\ , \\
D = 1 \quad &: \quad a = 0.21005\ , \quad b = 0.52762\ , \quad c = 1.14031\ , \quad R^2 = 0.99998\ , \\
D = 0.4 \quad &: \quad a = 0.31055\ , \quad b = 0.93641\ , \quad c = 1.79150\ , \quad R^2 = 0.99986\ , \\
D = 0.1 \quad &: \quad a = 0.58997\ , \quad b = 3.11843\ , \quad c = 4.16241\ , \quad R^2 = 0.99810\ , \\
D = 0.04 \quad &: \quad a = 1.12850\ , \quad b = 15.7413\ , \quad c = 10.6911\ , \quad R^2 = 0.99660\ , \\
D = 0.04/1.55 \quad &: \quad a = 2.33575\ , \quad b = 64.5058\ , \quad c = 18.8357\ , \quad R^2 = 0.99877.
\end{aligned}$$

Using the fitting function, we obtain

$$\int_{x_-/\sqrt{2D}}^{x_+/\sqrt{2D}} dz e^{z^2} \text{erfc}(z) = a + b \exp\left(-c\frac{x_- + x_+}{2}\right) \ , \tag{28}$$

$$\int_{y_-/\sqrt{2D}}^{y_+/\sqrt{2D}} dz e^{z^2} \text{erfc}(z) = a + b \exp\left(-c\frac{y_- + y_+}{2}\right) \ . \tag{29}$$

Substituting the absorbing and reset points $x_\pm$, $y_\pm$ Eq. (24) into the fitting functions in Eq. (28) and Eq. (29), we find the mean first passage time $\tau_{\text{off}}$:

$$\begin{aligned}
\tau_{\text{off}} &\approx \frac{\sqrt{\pi}}{1+g}\left[a + b\exp\left(-c[\frac{1}{1+g}h - \frac{f-g}{1+g}]\right)\right] \\
&= \frac{\sqrt{\pi}}{1+g}\left[a + b_1\exp\left(-\frac{c}{1+g}h\right)\right] \ ,
\end{aligned} \tag{30}$$

where the constant $b_1$ is defined as

$$b_1 = b \exp\left(c\frac{f-g}{1+g}\right) \ . \tag{31}$$

Similarly, the mean first passage time $\tau_{\text{on}}$ is:

$$\begin{aligned}
\tau_{\text{on}} &\approx \frac{\sqrt{\pi}}{1+g}\left[a + b\exp\left(-c[-\frac{1}{1+g}h + \frac{f+g}{1+g}]\right)\right] \\
&= \frac{\sqrt{\pi}}{1+g}\left[a + b_2\exp\left(\frac{c}{1+g}h\right)\right] \ ,
\end{aligned} \tag{32}$$

where $b_2$ is defined as

$$b_2 = b \exp\left(-c\frac{f+g}{1+g}\right) . \tag{33}$$

###### 2.2.5 Analytical reduction of the dynamical system model to a binary-unit network

Since in the bistable regime, each unit in the dynamical-system model can be approximated as a binary unit $S_i = \{0, 1\}$, we approximated the spatio-temporal dynamical system Eq. (14) by a network of binary units. We derived analytical expressions for parameters of the binary-unit network using parameters in the corresponding dynamical system model.

The reduced model consists of $N$ binary units $S_i$ ($i = 1, ..., N$), where $S_i = 1$ and $S_i = 0$ correspond to the On and Off phases, respectively. The dynamics of the binary-unit model is described by the transition rates $\omega(S_i = 0) = \omega(0 \to 1)$ and $\omega(S_i = 1) = \omega(1 \to 0)$. The transition rate $w(S_i)$ has the following properties:

1. $w(S_i)$ only depends on $S_i$ and its nearest neighbors, denoted as $S_{i\pm1}$. According to the corresponding dynamical system model, the index $i$ represents indices in two-dimensions $i = (x, y)$, and the interaction with the nearest neighbors has a discrete Laplacian form:

$$S_{i\pm1} = S_{x+1,y} - S_{x,y} + S_{x-1,y} - S_{x,y} + S_{x,y+1} - S_{x,y} + S_{x,y-1} - S_{x,y} . \tag{34}$$

2. When unit $i$ is in the On phase $S_i = 1$, its transition rate from $1$ to $0$ decreases if the nearest neighbor units are also in the On phase, which prolongs the On phase on average. The transition rate from $1$ to $0$ increases if the nearest neighbor units are in the Off phase.

3. When unit $i$ is in the Off phase $S_i = 0$, its transition rate from $0$ to $1$ increases if the nearest neighbor units are in the On phase. The transition rate from $0$ to $1$ decreases if the nearest neighbor units are in the Off phase.

The simplest form of a transition rate with these properties is:

$$w(S_i = 0) = \alpha_1 + \beta_1 S_{i\pm1}, \tag{35}$$

$$w(S_i = 1) = \alpha_2 - \beta_2 S_{i\pm1}, \tag{36}$$

where $\alpha_1, \alpha_2, \beta_1, \beta_2$ are positive numbers, and $S_{i\pm1}$ represents the nearest neighbour units of $S_i$. Using the property that $S_i = 0, 1$, a universal form of the transition rate can be written as

$$w(S_i) = w(S_i = 0) + [w(S_i = 1) - w(S_i = 0)]S_i . \tag{37}$$

Explicitly, we have

$$\begin{aligned} w(S_i) &= \alpha_1 + \beta_1 S_{i\pm1} + [\alpha_2 - \beta_2 S_{i\pm1} - \alpha_1 - \beta_1 S_{i\pm1}]S_i \\ &= \alpha_1 + (\alpha_2 - \alpha_1)S_i + \beta_1 S_{i\pm1} - (\beta_1 + \beta_2)S_i S_{i\pm1} . \end{aligned} \tag{38}$$

The transition rates $\omega(S_i = 0)$ and $\omega(S_i = 1)$ are the inverse of the mean On and Off episode durations, respectively. Therefore, we use Eq. (30) and Eq. (32) to derive expressions for $\omega(S_i = 0)$ and $\omega(S_i = 1)$ using parameters of the dynamical system model, and calculate $\alpha_1$, $\alpha_2$, $\beta_1$, $\beta_2$.

The mean episode durations Eq. (30) and Eq. (32) are functions of the intercept $h$, which contains the contributions from local recurrent interactions and the external input. We approximate the local interaction term $W\nabla^2 v$ as a function of binary variables $S_{i\pm1} \in \{0,1\}$. In the dynamical system model operating in the bistable regime, the average activity of a unit $i$ can be approximated as $v_{\text{on}}$ and $v_{\text{off}}$ during the On and Off phase, respectively. Here $v_{\text{on}}$ and $v_{\text{off}}$ are the values of $v$ at the two fixed points. Therefore, the activity of an arbitrary unit $i$ can be estimated as

$$v(i) \approx (v_{\text{on}} - v_{\text{off}})S_i + v_{\text{off}} \ . \tag{39}$$

The interaction term $W\nabla^2 v$ in the dynamical system Eq. (14) transforms into

$$\nabla^2 v \to \sum_{i\pm1} v(i \pm 1) \approx (v_{\text{on}} - v_{\text{off}})S_{i\pm1} \ , \tag{40}$$

where $S_{i\pm1}$ is the discrete Laplacian Eq. (34).

The values of $v_{\text{on}}$ and $v_{\text{off}}$ are given by the solution of the deterministic equations:

$$\begin{cases} \mp 1 - v - u + h = 0 \ , \\ gv - u + f = 0 \ , \end{cases} \tag{41}$$

$$v_{\text{on}} = \frac{1 + h - f}{1 + g} \ , \quad v_{\text{off}} = \frac{-1 + h - f}{1 + g} \ . \tag{42}$$

The intercept $h$ can be estimated as

$$h = W\nabla^2 v + I \approx W[(v_{\text{on}} - v_{\text{off}})S_{i\pm1}] + I \ . \tag{43}$$

From Eq. (42), we find $(v_{\text{on}} - v_{\text{off}}) = 2/(1 + g)$, so that $h$ can be written as

$$h \approx W\left[\frac{2}{1+g} S_{i\pm1}\right] + I \ . \tag{44}$$

Substituting this expression for $h$ into Eq. (30) and Eq. (32), we obtain $\tau_{\text{off}}$ and $\tau_{\text{on}}$ as functions of $I$ and $S_{i\pm1}$. For convenience, we define the positive constants

$$\begin{aligned} A_1 &= \frac{\sqrt{\pi}}{1+g}a \,, \\ B_1 &= \frac{\sqrt{\pi}}{1+g}b_1 \,, \\ B_2 &= \frac{\sqrt{\pi}}{1+g}b_2 \,, \\ D_1 &= \frac{c}{1+g} \cdot \frac{2W}{1+g} \,, \\ C_1 &= \frac{c}{1+g} \,. \end{aligned} \tag{45}$$

The explicit analytical expression for the transition rate $w(S_i = 0)$ is then given by

$$w(S_i = 0) = \frac{1}{\tau_{\text{off}}} = \frac{1}{A_1 + B_1 \exp[-D_1 S_{i\pm1} - C_1 I]}. \tag{46}$$

This equation shows that mean Off episode duration exponentially decreases with the external input current $\tau_{\text{off}} \sim \exp[-C_1 I]$, which is confirmed by numerical simulations. The term $\exp[-D_1(S_{\pm1})]$ captures in the effect of local recurrent interactions on the transition rate. The magnitude of this term is controlled by $D_1$, which is proportional to the interaction strength $W$. Assuming the interaction strength is relatively weak, we perform the Taylor expansion with respect to $D_1$ and retain the leading interaction term $D_1 S_{i\pm1}$. In this first-order approximation, the transition rate $w(S_i = 0)$ is expressed as

$$\begin{aligned} w(S_i = 0) &\approx \frac{1}{A_1 + B_1 \exp[-C_1 I][1 - D_1 S_{i\pm1}]} \\ &\approx \frac{1}{A_1 + B_1 \exp[-C_1 I]} + \frac{B_1 D_1 \exp[-C_1 I]}{(A_1 + B_1 \exp[-C_1 I])^2} S_{i\pm1}. \end{aligned}$$

From this equation, we find the parameters $\alpha_1$ and $\beta_1$ of the Off-to-On transition rate Eq. (35):

$$\alpha_1 = \frac{1}{A_1 + B_1 \exp[-C_1 I]}, \tag{47}$$

$$\beta_1 = \frac{B_1 D_1 \exp[-C_1 I]}{(A_1 + B_1 \exp[-C_1 I])^2}. \tag{48}$$

The values of $\alpha_1$ and $\beta_1$ depend on the input current $I$ and other parameters of the dynamical system model. The baseline rate $\alpha_1$ is the dominant term in the expression for the Off-to-On transition rate. In the limit of vanishing interactions $\tau_{\text{off}} \to 1/\alpha_1$. The rate $\alpha_1$ is a monotonically increasing function of input $I$. It has the minimum value $1/(A_1 + B_1)$ when $I = 0$, and it approaches $1/A_1$ when $I \to \infty$. The coupling strength $\beta_1$ characterizes the leading term of interactions between neighbouring units $S_{i\pm1}$. It is proportional to $D_1$, hence to the interaction strength parameter $W$. Its magnitude also depends on the input $I$, and it is a decreasing function of $I$ when

$$I \geqslant -\frac{\ln(A_1/B_1)}{C_1} = -\frac{\ln(a/b_1)}{C_1} = -\frac{1+g}{c}\left[\ln\left(\frac{a}{b}\right) - c\frac{f-g}{1+g}\right] . \quad (49)$$

Similarly, the analytic expression for the On-to-Off transition rate $w(S_i = 1)$ is

$$w(S_i = 1) = \frac{1}{\tau_{\text{on}}} = \frac{1}{A_1 + B_2 \exp[D_1 S_{i\pm1} + C_1 I]}. \quad (50)$$

This equation shows that the mean On episode duration exponentially increases with the external input current $\tau_{\text{on}} \sim \exp[C_1(I_{\text{stim}} + I_{\text{attn}})]$, which agrees with simulations. Applying Taylor expansion and keeping only the leading term of $S_{\pm1}$, we find

$$\begin{aligned} w(S_i = 1) &\approx \frac{1}{A_1 + B_2 \exp[C_1 I][1 + D_1 S_{i\pm1}]} \\ &\approx \frac{1}{A_1 + B_2 \exp[C_1 I]} - \frac{B_2 D_1 \exp[-C_1 I]}{(A_1 \exp[-C_1 I] + B_2)^2} S_{i\pm1}. \end{aligned}$$

From this equation, we find the parameters $\alpha_2$ and $\beta_2$ of the On-to-Off transition rate Eq. (36):

$$\alpha_2 = \frac{1}{A_1 + B_2 \exp[C_1 I]}, \quad (51)$$

$$\beta_2 = \frac{B_2 D_1 \exp[-C_1 I]}{(A_1 \exp[-C_1 I] + B_2)^2}, \quad (52)$$

where $\alpha_2$ and $\beta_2$ are positive. Comparing the expressions for $w(S_i = 0)$ Eq. (35) and $w(S_i = 1)$ Eq. (36), we see that instead of a "+" sign in front of $\beta_1$, we have a "$-$" sign in front of $\beta_2$. The "$-$" sign means that the recurrent interactions favor nearby units to stay in the same phases. The transition rates reduce when two nearby units are in the same phase, whereas the transition rates increase when nearby units are in different phases.

The values of $\alpha_2, \beta_2$ depend on the input current $I$ and other parameters of the dynamical system model. The baseline rate $\alpha_2$ is the dominant term in the On-to-Off transition rate. In the limit of vanishing interactions, $\tau_{\text{on}} \to 1/\alpha_2$. The rate $\alpha_2$ is a monotonically decreasing function of $I$. It has the maximum value $1/(A_1 + B_2)$ when $I = 0$, and it decreases exponentially and approaches zero when $I \to \infty$:

$$\alpha_2 \to \frac{1}{B_2} \exp[-C_1(I)]. \quad (53)$$

Thus, the average On episode duration increases with increasing external current and approaches infinity in the large current limit. As a result, the ratio $\alpha_2/\alpha_1$, or approximately $\tau_{\rm off}/\tau_{\rm on}$ is a monotonically decreasing function of $I$. Larger positive external input drives the units to stay in the On phase longer.

The coupling strength $\beta_2$ characterizes the leading term of interactions between neighbouring units $S_{i\pm1}$. It is proportional to $D_1$, hence to the interaction strength $W$, and its magnitude also depends on input $I$. It is a decreasing function of $I$ when

$$I \geqslant -\frac{\ln(B_2/A_1)}{C_1} = -\frac{\ln(b_2/a)}{C_1} = -\frac{1+g}{c}\left[\ln\left(\frac{b}{a}\right) - c\frac{f+g}{1+g}\right]. \tag{54}$$

In the large $I$ limit, $\beta_1$ decreases exponentially with increasing $I$:

$$\beta_2 \to \frac{D_1}{B_2}\exp[-C_1 I]. \tag{55}$$

Thus, we analytical reduced the dynamical system model of spatiotemporal On-Off dynamics to a binary-unit network, where the rate parameters $\alpha_1$, $\alpha_2$, $\beta_1$, and $\beta_2$ are expressed through the parameters of the dynamical system model. In the dynamical system model, we modeled attention effect by changing the external input current $I$, which corresponds to changes of the rate parameters $\alpha_1$, $\alpha_2$, $\beta_1$ and $\beta_2$ in the reduced binary-unit model.

#### 2.3 Analytical calculation of noise correlations in the binary-unit network

We used the reduced binary-unit network model to analytical derive the dependence of noise correlations on the lateral distance and attention. In this reduced model, each unit is described by a binary variable $S_i = \{0, 1\}$, $(i = 1, ..., N)$. At time $t$, the probability of the network to be in a certain configuration $\{S\} = \{S_1, S_2, ..., S_N\}$ is denoted as $P(\{S\}, t)$. The time evolution of $P(\{S\}, t)$ is described by the master equation:

$$\frac{d}{dt}P(\{S\}, t) = -P(\{S\}, t)\sum_i w(S_i) + \sum_i P(\{S\}^{i*}, t)w(1 - S_i). \tag{56}$$

Here $\{S\}^{i*} = \{S_1, S_2, ..., 1 - S_i, ..., S_N\}$, and $w(S_i)$ are the transition rates.

##### 2.3.1 Moment expansion

Using the master equation, we can write the equations for time evolution of arbitrary moments of $S_i$. For example, the average activity of units $S_i$ is defined as

$$\langle S_i\rangle(t) = \sum_{\{S\}} P(\{S\}, t)S_i\ , \tag{57}$$

where the summation is over all configurations $\{S\}$ at time $t$. Its time evolution is given by

$$\frac{d}{dt}\langle S_i\rangle(t) = \frac{d}{dt}\left(\sum_{\{S\}} P(\{S\},t)S_i\right) = \sum_{\{S\}}\left(\frac{d}{dt}P(\{S\},t)\right)S_i \; . \tag{58}$$

Substituting the master equation, we find

$$\frac{d}{dt}\langle S_i\rangle(t) = \sum_{\{S\}} P(\{S\},t)[w(S_i)(1-2S_i)] \; . \tag{59}$$

Similarly, the rate of change of average activity of a pair of units is

$$\frac{d}{dt}\langle S_iS_j\rangle(t) = \sum_{\{S\}} P(\{S\},t)[w(S_i)(1-2S_i)S_j + w(S_j)(1-2S_j)S_i] \; . \tag{60}$$

Substituting the transition rates in Eq. (38) and summing over all configurations, we get the coupled equations for moments [53]:

$$\frac{d}{dt}\langle S_i\rangle(t) = \alpha_1 - (\alpha_1+\alpha_2)\langle S_i\rangle + \beta_1\langle S_{i\pm1}\rangle + (\beta_2-\beta_1)\langle S_iS_{i\pm1}\rangle \; , \tag{61}$$

and

$$\begin{aligned}\frac{d}{dt}\langle S_iS_j\rangle(t) &= \alpha_1(\langle S_i\rangle + \langle S_j\rangle) - 2(\alpha_1+\alpha_2)\langle S_iS_j\rangle + \beta_1(\langle S_{i\pm1}S_j\rangle + \langle S_{j\pm1}S_i\rangle) \\ &+ (\beta_2-\beta_1)(\langle S_iS_{i\pm1}S_j\rangle + \langle S_jS_{j\pm1}S_i\rangle) \; , \; (i\neq j) \; .\end{aligned} \tag{62}$$

Here we used the property of binary variables $S_i = S_i^2$ to simplify the equations. The system of equations Eq. (61) and Eq. (62) is not closed since it involve moments of cubic order($\langle SSS\rangle$), therefore, it cannot be solved in general. However, since the fluctuation of activity $\delta S \ll 1$, the higher order moments are suppressed. Therefore, as an approximation, we neglect the cubic and higher moments to simplify the equations:

$$\begin{aligned}\frac{d}{dt}\langle S_i\rangle(t) &= \alpha_1 - (\alpha_1+\alpha_2)\langle S_i\rangle + \beta_1\langle S_{i\pm1}\rangle \; , \\ \frac{d}{dt}\langle S_iS_j\rangle(t) &= \alpha_1(\langle S_i\rangle + \langle S_j\rangle) - 2(\alpha_1+\alpha_2)\langle S_iS_j\rangle + \beta_1(\langle S_{i\pm1}S_j\rangle + \langle S_{j\pm1}S_i\rangle) \; .\end{aligned} \tag{63}$$

Since we are interested in the population average of the first moment $\langle S\rangle = (\sum_i \langle S_i\rangle)/N$, we sum over the index $i$ and take the average of the first equation in Eqs. (63):

$$\frac{d}{dt}\langle S\rangle(t) = \alpha_1 - (\alpha_1+\alpha_2)\langle S\rangle \; . \tag{64}$$

The solution of this equation is given by

$$\langle S\rangle(t) = [S(0) - S(\infty)]\exp(-\lambda_0 t) + S(\infty) \;, \tag{65}$$

where $\lambda_0$ is

$$\lambda_0 = \alpha_1 + \alpha_2 \;, \tag{66}$$

and the steady state at infinite time is

$$\langle S\rangle(t \to \infty) = S(\infty) = \frac{\alpha_1}{\alpha_1 + \alpha_2} \;. \tag{67}$$

##### 2.3.2 Time-delayed correlation functions

To calculate noise correlation $r_{\mathrm{sc}}$, we need explicit expressions for time-delayed correlation functions. The time evolution of the time-delayed quadratic moment is defined as [53]

$$\frac{d}{d\tau}\langle S_i(t)S_j(t+\tau)\rangle = \sum_{\{S\}} P(\{S\},t) S_i \frac{d}{d\tau}\left(\sum_{\{\sigma\}} P(\{\sigma\},t+\tau|\{S\},t)\sigma_j\right) \;. \tag{68}$$

Here $P(\{\sigma\},t+\tau|\{S\},t)$ is the conditional probability, which is the probability of finding the system in the configuration $\{\sigma\}$ at time $t+\tau$, given that it was in the configuration $\{S\}$ at time $t$. Since the conditional probability obeys the same master equation as in Eq. (56), we have

$$\begin{aligned} \frac{d}{d\tau}\langle S_i(t)S_j(t+\tau)\rangle &= \langle S_i(t)(1-2S_j(t+\tau))w(S_j(t+\tau))\rangle \\ &= \alpha_1\langle S_i(t)\rangle - (\alpha_1+\alpha_2)\langle S_i(t)S_j(t+\tau)\rangle + \beta_1\langle S_i(t)S_{j\pm 1}(t+\tau)\end{aligned} \tag{69}$$

In our analysis, we are interested in the equilibrium value of the time-delayed quadratic moment, which amounts to taking the limit $t \to \infty$ while keeping $\tau$ finite. We define $\lim_{t\to\infty}\langle S_i(t)S_j(t+\tau)\rangle = G_{ij}(\tau)$ and obtain

$$\frac{d}{d\tau}G_{ij}(\tau) = \alpha_1\langle S_i(\infty)\rangle - (\alpha_1+\alpha_2)G_{ij}(\tau) + \beta_1 G_{ij\pm 1}(\tau) \;. \tag{70}$$

In the continuum limit, Eq. (70) becomes

$$\frac{d}{d\tau}G_{ij}(\tau) = \alpha_1\langle S_i(\infty)\rangle - (\alpha_1+\alpha_2)G_{ij}(\tau) + \beta_1[(\Delta d)^2\nabla^2 G_{ij}(\tau)] \;. \tag{71}$$

To evaluate the distance-dependence of noise correlations, we need to compute the averaged quantity $G(d,\tau) = 1/N_d \sum_{|i-j|=d} G_{ij}(\tau)$, where $N_d$ is the number of unit pairs with the distance $d$. This quantity only depends on the absolute values $|i-j| = d$ and $|\tau|$, so it is symmetric under the exchange of indices $i$ and $j$. For the equal-time quadratic moment $\langle S_i(t)S_j(t)\rangle$, this averaging is straightforward. However, for the time-delayed quadratic moment $\langle S_i(t)S_j(t+\tau)\rangle$, the indices $i$ and $j$ are not symmetric, which needs to be taken into account in averaging. First, on the left-hand side of Eq. (71), the operator $d/d\tau$ only acts on the variable with the index $j$, $S_j(t+\tau)$. Averaging both sides of Eq. (71) results in $G(d,\tau)$ that is symmetric in $i \leftrightarrow j$, so that the variable with index $i$ also has the same $\tau$ dependence as that with $j$. Since the operator $d/d\tau$ in Eq. (71) is asymmetric and only acts on $j$, when it acts on $G(d,\tau)$, it only contains half of the action of time derivative. Therefore, we need to add a factor $1/2$ in the replacement $\frac{d}{d\tau}G_{ij}(\tau) \to \frac{1}{2}\frac{d}{d\tau}G(d,\tau)$. Second, on the right-hand side of Eq. (71), we use the continuum limit approximation $G_{ij\pm 1}(\tau) \approx (\Delta d)^2\nabla^2 G_{ij}(\tau)$. In this expression, the term $\nabla^2 G_{ij}(\tau)$ accounts for changing the index $j$, namely, $\Delta d \propto |j \pm 1 - i|$. Again, since after averaging $G(d,\tau)$ is symmetric in $i$ and $j$, the operator $\nabla^2 G(d,\tau)$ contains terms induced by changing indices of both $i$ and $j$, therefore we need to include a factor $1/2$ in the replacement $\nabla^2 G_{ij}(\tau) \to \frac{1}{2}\nabla^2 G(d,\tau)$. Taking these factors into account, we obtain the equation for time evolution of the averaged time-delayed quadratic moment $G(d,\tau)$:

$$\frac{1}{2}\frac{d}{d\tau}G(d,\tau) = \alpha_1 S(\infty) - (\alpha_1+\alpha_2)G(d,\tau) + \frac{1}{2}\beta_1(\Delta d)^2\nabla^2 G(d,\tau) \,. \tag{72}$$

The physical solution of this equation is given by

$$G(d,\tau) = [S(\infty)]^2 + [1-S(\infty)]S(\infty)\exp\left(-\frac{d}{L} - \frac{|\tau|}{\tau_c}\right) \,, \quad d = |i-j| \,, \tag{73}$$

where the correlation length $L$ is

$$L = \sqrt{\frac{\beta_1}{\alpha_1+\alpha_2}} \,, \tag{74}$$

and the time constant $\tau_c$ is given by

$$\tau_c = \frac{1}{\lambda_0} = \frac{1}{\alpha_1+\alpha_2} \,. \tag{75}$$

In the limit $d \to 0$, $G(d,\tau)$ reduces to the auto-correlation function:

$$G(0,\tau) = [S(\infty)]^2 + [1-S(\infty)]S(\infty)\exp\left(-\frac{|\tau|}{\tau_c}\right) \,. \tag{76}$$

In the limit of no interactions $\beta_1 \to 0$, $G(0,\tau)$ reduces to

$$G(0,\tau|\beta_1 \to 0) = \left(\frac{\alpha_1}{\alpha_1+\alpha_2}\right)^2 + \frac{\alpha_1\alpha_2}{(\alpha_1+\alpha_2)^2}\exp\left(-(\alpha_1+\alpha_2)|\tau|\right) \,. \tag{77}$$

If we use the notation

$$\alpha_1 = \frac{1}{\tau_{\text{off}}} \,, \qquad \alpha_2 = \frac{1}{\tau_{\text{on}}} \,, \quad (\beta_1 = 0) \,, \tag{78}$$

we obtain the following expression for the auto-correlation:

$$G(0,\tau|\beta_1 \to 0) = \left(\frac{\tau_{\text{on}}}{\tau_{\text{off}}+\tau_{\text{on}}}\right)^2 + \frac{\tau_{\text{on}}\tau_{\text{off}}}{(\tau_{\text{on}}+\tau_{\text{off}})^2}\exp\left(-\frac{\tau_{\text{on}}+\tau_{\text{off}}}{\tau_{\text{on}}\tau_{\text{off}}}|\tau|\right) \ , \tag{79}$$

which correctly matches the steady-state value of the time-delayed quadratic moment in single-column On-Off dynamics without lateral interactions.

##### 2.3.3 Noise correlations

We derived analytical expressions for noise correlations $r_{\text{sc}}$ in the binary-unit network model. We label neurons with indices $(\mathbf{x}, i)$, where $\mathbf{x}$ is the index of 2-dimensional lateral position along cortical surface and $i$ indexes neurons within the same column (i.e. along laminar direction perpendicular to cortical surface). All neurons $(\mathbf{x}, i)(i = 1, 2...)$ with the shared lateral index $\mathbf{x}$ switch between the On and Off phases via a Markov process, which is described by a single binary variable $S(\mathbf{x}, t) = \{0, 1\}$, where $S(\mathbf{x}, t) = 1$ and $S(\mathbf{x}, t) = 0$ denote the On and Off phases, respectively. In particular, $S(\mathbf{x}, t)$ is independent of index $i$. In other words, the On-Off dynamics of all neurons are synchronous within a cortical column with index $\mathbf{x}$. The mean firing rate during the On and Off phases are $r_{\text{on}}(\mathbf{x}, i)$ and $r_{\text{off}}(\mathbf{x}, i)$, respectively. The mean firing rates also depend on the neuron index $i$ within a column.

With this notation, the mean firing rate as a function of time $t$ is

$$\lambda(\mathbf{x}, i; t) = r_{\text{off}}(\mathbf{x}, i) + S(\mathbf{x}, t)[r_{\text{on}}(\mathbf{x}, i) - r_{\text{off}}(\mathbf{x}, i)] \ . \tag{80}$$

The time-integral of this firing rate over the measurement time-window $T$ is

$$\Lambda(\mathbf{x}, i) = \int_t^{t+T} \lambda(\mathbf{x}, i; t')dt' = r_{\text{off}}(\mathbf{x}, i)T + R(\mathbf{x})\Delta r(\mathbf{x}, i) \ . \tag{81}$$

Here $\Delta r(\mathbf{x}, i)$ is defined as

$$\Delta r(\mathbf{x}, i) = r_{\text{on}}(\mathbf{x}, i) - r_{\text{off}}(\mathbf{x}, i) \ , \tag{82}$$

and the normalized rate $R(\mathbf{x})$ is

$$R(\mathbf{x}) = \int_t^{t+T} S(\mathbf{x}, t')dt' \ . \tag{83}$$

We assume that neurons emit spikes as inhomogeneous Poisson processes, so the number of spike $N(\mathbf{x}, i)$ produced on each trial obeys the Poisson distribution:

$$P(N(\mathbf{x}, i) = n) = \frac{(\Lambda(\mathbf{x}, i))^n}{n!} e^{-\Lambda(\mathbf{x}, i)} \ . \tag{84}$$

The spike count $N(\mathbf{x}, i)$ depends on the binary variable $S(\mathbf{x}, t)$ via the normalized rate $R(\mathbf{x}, t)$, which fluctuates from trial to trial. To calculate the spike-count correlations, we need to calculate moments of the spike-count averaged over trials. For sufficient number of trials, the trial-average includes the average of binary variable $S(\mathbf{x}, t)$ according to the probability defined in the master equation, $P(\{S\}, t)$. Here we use the symbol $\langle\ \rangle$ to represent this average.

The mean value of spike-count $N(\mathbf{x}, i)$ is given by the double average over trials and the Poisson distribution:

$$\mathrm{E}[N(\mathbf{x}, i)] = \langle \sum_{n=1}^{\infty} n \frac{(\Lambda(\mathbf{x}, i))^n}{n!} e^{-\Lambda(\mathbf{x},i)} \rangle = \langle \Lambda(\mathbf{x}, i) \rangle = r_{\text{off}}(\mathbf{x}, i)T + \mathrm{E}[R(\mathbf{x})]\Delta r(\mathbf{x}, i) \ . \quad (85)$$

Similarly, the variance of the spike-count $N(\mathbf{x}, i)$ is given by

$$\begin{aligned} \mathrm{Var}[N(\mathbf{x}, i)] &= \mathrm{E}[N(\mathbf{x}, i)^2] - (\mathrm{E}[N(\mathbf{x}, i)])^2 = \langle \sum_{n=1}^{\infty} n^2 \frac{(\Lambda(\mathbf{x}, i))^n}{n!} e^{-\Lambda(\mathbf{x},i)} \rangle - (\langle \Lambda(\mathbf{x}, i) \rangle)^2 \\ &= \langle [\Lambda(\mathbf{x}, i)]^2 + \Lambda(\mathbf{x}, i) \rangle - (\langle \Lambda(\mathbf{x}, i) \rangle)^2 \\ &= (\Delta r(\mathbf{x}, i))^2 \mathrm{Var}[R(\mathbf{x})] + r_{\text{off}}(\mathbf{x}, i)T + \mathrm{E}[R(\mathbf{x})]\Delta r(\mathbf{x}, i) \ . \end{aligned} \quad (86)$$

The covariance of spike-counts of a pair of neurons $\mathrm{Cov}[N(\mathbf{x}, i), N(\mathbf{y}, j)]$ is

$$\begin{aligned} \mathrm{Cov}[N(\mathbf{x}, i), N(\mathbf{y}, j)] &= \mathrm{E}[N(\mathbf{x}, i)N(\mathbf{y}, j)] - \mathrm{E}[N(\mathbf{x}, i)]\mathrm{E}[N(\mathbf{y}, j)] \\ &= \langle \sum_{n=1}^{\infty} n \frac{(\Lambda(\mathbf{x}, i))^n}{n!} e^{-\Lambda(\mathbf{x},i)} \sum_{m=1}^{\infty} m \frac{(\Lambda(\mathbf{y}, j))^m}{m!} e^{-\Lambda(\mathbf{y},j)} \rangle - \langle \Lambda(\mathbf{x}, i) \rangle \langle \Lambda(\mathbf{y}, j) \rangle \\ &= \langle \Lambda(\mathbf{x}, i)\Lambda(\mathbf{y}, j) \rangle - \langle \Lambda(\mathbf{x}, i) \rangle \langle \Lambda(\mathbf{y}, j) \rangle \\ &= \Delta r(\mathbf{x}, i)\Delta r(\mathbf{y}, j)\mathrm{Cov}[R(\mathbf{x}), R(\mathbf{y})] \ . \end{aligned} \quad (87)$$

In this equation, we notice that if $\mathbf{x} = \mathbf{y}$, we obtain

$$\mathrm{Cov}[N(\mathbf{x}, i), N(\mathbf{x}, j)] = \Delta r(\mathbf{x}, i)\Delta r(\mathbf{x}, j)\mathrm{Var}[R(\mathbf{x})] \ , \quad (88)$$

which is the same as Eq. (9) we obtained earlier for a single column.

Using these expressions for $\mathrm{Var}[N(\mathbf{x}, i)]$ and $\mathrm{Cov}[N(\mathbf{x}, i), N(\mathbf{y}, j)]$, we can calculate the noise correlation $r_{\text{sc}}$. For a pair of neurons with indices $(\mathbf{x}, i)$ and $(\mathbf{y}, j)$, the noise correlation is

$$r_{\text{sc}} = \frac{\mathrm{Cov}[N(\mathbf{x}, i), N(\mathbf{y}, j)]}{\sqrt{\mathrm{Var}[N(\mathbf{x}, i)]\mathrm{Var}[N(\mathbf{y}, j)]}} = \frac{\Delta r(\mathbf{x}, i)\Delta r(\mathbf{y}, j)\mathrm{Cov}[R(\mathbf{x}), R(\mathbf{y})]}{\sqrt{\mathrm{Var}[N(\mathbf{x}, i)]\mathrm{Var}[N(\mathbf{y}, j)]}} \ . \quad (89)$$

In the special case where two neurons are in the same column, namely $\mathbf{x} = \mathbf{y}$, this expression reduces to

$$r_{\text{sc}} = \frac{\Delta r(\mathbf{x}, i)\Delta r(\mathbf{x}, j)\mathrm{Var}[R(\mathbf{x})]}{\sqrt{\mathrm{Var}[N(\mathbf{x}, i)]\mathrm{Var}[N(\mathbf{x}, j)]}} \ , \quad (90)$$

which is the same as the expression for noise correlations within a single column derived in section 2.1.1.

Noise correlations depend on two sets of parameters. The first set is directly extracted from the data: time window $T$ and mean firing rates during the On and Off phases $r_{\text{off}}(\mathbf{x}, i)$, $r_{\text{on}}(\mathbf{x}, i)$. We estimate these parameters from HMM fitting of local columnar On-Off dynamics. The second set contains statistical quantities of On-Off dynamics, namely, $\text{E}[R(\mathbf{x})]$, $\text{Var}[R(\mathbf{x})]$ and $\text{Cov}[R(\mathbf{x}), R(\mathbf{y})]$. These statistical quantities are calculated based on the model of On-Off dynamics and expressed as functions of the model parameters, whcih are also estimated by HMM. In the calculation, we used the steady-state solution of the model ($t = \infty$).

The mean of the normalized rate $\text{E}[R(\mathbf{x})]$ is a time integral of the mean of the binary variable $S(\mathbf{x}, t)$, which is

$$\text{E}[R(\mathbf{x})] = \int_t^{t+T} \langle S(\mathbf{x}, t') \rangle dt' = \int_t^{t+T} \langle S(\mathbf{x}, t' \to \infty) \rangle dt' = S(\infty) T = \frac{\alpha_1}{\alpha_1 + \alpha_2} T \; . \quad (91)$$

In the limit of no interactions between nearby units $\beta_1 \to 0$, we obtain

$$\text{E}[R(\mathbf{x})] = \frac{\alpha_1}{\alpha_1 + \alpha_2} T = \frac{\tau_{\text{on}}}{\tau_{\text{on}} + \tau_{\text{off}}} T \; , \quad \left( \alpha_1 = \frac{1}{\tau_{\text{off}}} \; , \; \alpha_2 = \frac{1}{\tau_{\text{on}}} \right) \; , \quad (92)$$

which is the same as Eq. (10) derived earlier for single columns.

The covariance of normalized rates Cov[R($\mathbf{x}$),R($\mathbf{y}$)] is

$$\text{Cov}[R(\mathbf{x}), R(\mathbf{y})] = \langle \int_0^T S(\mathbf{x}, t_1) dt_1 \int_0^T S(\mathbf{y}, t_2) dt_2 \rangle - \text{E}[R(\mathbf{x})] \cdot \text{E}[R(\mathbf{y})] \; , \quad (93)$$

where, for convenience, we shifted the limits of time integral from $[t, t+T]$ to $[0, T]$. The first term in the covariance involves a double integral of the quadratic moment of the binary variable, which we convert to a double integral of the averaged time-delayed quadratic moment $G(|\mathbf{x} - \mathbf{y}|, \tau)$:

$$\begin{aligned} &\langle \int_0^T S(\mathbf{x}, t_1) dt_1 \int_0^T S(\mathbf{y}, t_2) dt_2 \rangle = \int_0^T dt_1 \int_0^T dt_2 \langle S(\mathbf{x}, t_1) S(\mathbf{y}, t_2) \rangle \\ &= 2 \int_0^T dt_1 \int_0^{t_1} dt_2 \langle S(\mathbf{x}, t_1) S(\mathbf{y}, t_2) \rangle \\ &= 2 \int_0^T dt_1 \int_0^{t_1} d\tau \langle S(\mathbf{x}, t_1) S(\mathbf{y}, t_1 - \tau) \rangle \\ &= 2 \int_0^T dt_1 \int_0^{t_1} d\tau G(|\mathbf{x} - \mathbf{y}|, \tau) \; . \end{aligned} \quad (94)$$

On the second line of Eq. (94), we used the property that the integrand $\langle S(\mathbf{x}, t_1) S(\mathbf{y}, t_2) \rangle$ is symmetric about the line $t_1 = t_2$, so the double integral $\int_0^T dt_1 \int_0^T dt_2$ is just twice the integral

$\int_0^T dt_1 \int_0^{t_1} dt_2$, resulting in the factor of 2. On the fourth line of Eq. (94), we used the fact that in the steady-state, $\langle S(\mathbf{x},t)S(\mathbf{y},t+\tau)\rangle$ only depends on $\tau$ and $|\mathbf{x}-\mathbf{y}|$, and $G(|\mathbf{x}-\mathbf{y}|,\tau)$ is an even function of $\tau$. Here the trial-average $\langle S(\mathbf{x},t)S(\mathbf{y},t+\tau)\rangle$ also includes the average of pairs of neurons with the same spatial separation $|\mathbf{x}-\mathbf{y}|$, so we have $\langle S(\mathbf{x},t)S(\mathbf{y},t-\tau)\rangle \to G(|\mathbf{x}-\mathbf{y}|,\tau)$.

With these transformations, we formally express the covariance as

$$\text{Cov}[R(\mathbf{x}),R(\mathbf{y})] = 2\left[\int_0^T dt_1 \int_0^{t_1} G(|\mathbf{x}-\mathbf{y}|,\tau)d\tau\right] - [S(\infty)]^2T^2\,, \tag{95}$$

and the variance as

$$\text{Var}[R(\mathbf{x})] = 2\left[\int_0^T dt_1 \int_0^{t_1} G(0,\tau)d\tau\right] - [S(\infty)]^2T^2\,. \tag{96}$$

These expressions for $\text{Cov}[R(\mathbf{x}),R(\mathbf{y})]$ and $\text{Var}[R(\mathbf{x})]$ are general and do not depend on a specific model of On-Off dynamics. Therefore, we can use Eq. (91), Eq. (95) and Eq. (96) to obtain an explicit expression of noise correlations for binary-unit network models with any types of interactions, once we calculate the time-delayed correlation $G(|\mathbf{x}-\mathbf{y}|,\tau)$ and the steady-state mean activity $S(\infty)$.

We apply Eq. (95) and Eq. (96) to our binary-unit network model. Performing the double time integrals, we get

$$\begin{aligned}\text{Cov}[R(\mathbf{x}),R(\mathbf{y})] &= 2\int_0^T dt_1 \int_0^{t_1}\left[[S(\infty)]^2 + [1-S(\infty)]S(\infty)\exp(-\frac{|\mathbf{x}-\mathbf{y}|}{L} - \frac{|\tau|}{\tau_c})\right]d\tau \\ &- [S(\infty)]^2T^2 \\ &= 2[1-S(\infty)]S(\infty)\exp(-\frac{|\mathbf{x}-\mathbf{y}|}{L})\tau_c\left[T - \tau_c\left(1-\exp\left(-\frac{T}{\tau_c}\right)\right)\right]\,,\end{aligned} \tag{97}$$

and

$$\text{Var}[R(\mathbf{x})] = 2[1-S(\infty)]S(\infty)\tau_c\left[T - \tau_c\left(1-\exp\left(-\frac{T}{\tau_c}\right)\right)\right]\,. \tag{98}$$

Substituting the explicit expressions in terms of the model parameters $\alpha_1, \alpha_2$ and $\beta_1$, we have

$$\begin{aligned}&\text{Cov}[R(\mathbf{x}),R(\mathbf{y})] = \\ &\frac{2\alpha_1\alpha_2}{(\alpha_1+\alpha_2)^3}\exp\left(-\frac{|\mathbf{x}-\mathbf{y}|}{L}\right)\left[T - \frac{1}{\alpha_1+\alpha_2}\left(1-\exp\left(-(\alpha_1+\alpha_2)T\right)\right)\right]\,,\end{aligned} \tag{99}$$

and

$$\text{Var}[R(\mathbf{x})] = \frac{2\alpha_1\alpha_2}{(\alpha_1+\alpha_2)^3}\left[T - \frac{1}{\alpha_1+\alpha_2}\left(1-\exp\left(-(\alpha_1+\alpha_2)T\right)\right)\right]\,. \tag{100}$$

In summary, we derived a general analytical expression Eq. (89) for the noise correlation $r_{\text{sc}}$ based on spatiotemporal On-Off dynamics. This equation describes noise correlations for pairs of neurons within the same column and in different columns. When two neurons are in different columns with lateral separation $|\mathbf{x}-\mathbf{y}|$, we find that the noise correlation is proportional to $\text{Cov}[R(\mathbf{x}), R(\mathbf{y})]$, which decays exponentially as a function of lateral distance, with the correlation length $L$:

$$r_{\text{sc}} \propto \text{Cov}[R(\mathbf{x}), R(\mathbf{y})] = \mathcal{A}(\alpha_1, \alpha_2) \exp\left(-\frac{|\mathbf{x}-\mathbf{y}|}{L}\right) . \tag{101}$$

Here $\mathcal{A}(\alpha_1, \alpha_2)$ describes noise correlations of two neurons within a single column, Eq. (6). The covariance $\text{Cov}[R(\mathbf{x}), R(\mathbf{y})]$ quantifies the coherence of On-Off dynamics for pairs of neurons located in different columns $\mathbf{x}$ and $\mathbf{y}$. The covariance vanishes when two neurons are far away from each other $|\mathbf{x}-\mathbf{y}| \gg L$. The exponential decay factor $\exp(-|\mathbf{x}-\mathbf{y}|/L)$ agrees with experimental observations that the average noise correlations decrease with lateral separation.

### 3 Spatial patterns of correlations in alternative models of network dynamics

We analyzed the spatial patterns of neural correlations arising from different types of network dynamics, which included a network model with local metastability (similar to our binary-unit network), the network with chaotic instability[24], and the network with fluctuations around a single stable fixed point induced by external noise[25]. We studied the spatial profile of noise correlations and its dependence on inputs, which can characterize patterns of correlations at different behavioral states, such as spontaneous activity or evoked activity during stimulus onset and attention. The spatial patterns of noise correlations provide us a window to infer the underlying dynamical mechanism.

#### 3.1 General propriety of neural correlations

We consider a firing rate model of a group of units $v_i$ ($i = 1....N$) with the dynamical equations given by

$$\frac{d}{dt} v_i(t) = -v_i + f(W_{ij} v_j + I) + \eta_i(t) . \tag{102}$$

Here $\eta_i(t)$ is a correlated external noisy input, which satisfies $\langle \eta_k(t) \eta_j(t+\tau) \rangle = [\mathbf{\Sigma}_0]_{ij} \exp(-\tau/\tau_0)$. The activity of units is assumed to fluctuate around a fixed point:

$$\bar{v}_i = f(W_{ij} \bar{v}_j + I) . \tag{103}$$

We define the fluctuation $\delta v_i = v_i - \bar{v}_i$ and study the linearized dynamics around the fixed point

$$\begin{aligned}\frac{d}{dt}\delta v_i(t) &= -\delta v_i + [f'(W_{ij}\bar{v}_j + I)W_{ij}]\delta v_j + \eta_i(t) \\ &= -\delta v_i + [W_{ij}^{\text{eff}}]\delta v_j + \eta_i(t) \; . \end{aligned} \tag{104}$$

Here we defined the effective interaction strength to be the product of the first-order derivative $f'$ and the connectivity $W_{ij}$, i.e. $W_{ij}^{\text{eff}} = f'(W_{ij}\bar{v}_j + I)W_{ij}$, which is a function of the external current $I$. In the matrix form, we have

$$\dot{\delta \mathbf{v}} = -(\mathbf{I} - \mathbf{W}^{\text{eff}})\delta\mathbf{v} + \eta(t) \; . \tag{105}$$

We can then derive the evolution equation of the time-delay correlation function:

$$\frac{d}{d\tau}\langle \delta v_i(t)\delta v_j(t+\tau)\rangle = -\langle \delta v_i(\mathbf{I} - \mathbf{W}^{\text{eff}})_{jk}\delta v_k\rangle + \langle \delta v_i(t)\eta_j(t+\tau)\rangle \; . \tag{106}$$

If we separate the autocorrelation $\langle \delta v_i(t)\delta v_i(t+\tau)\rangle = \mathcal{A}(\tau)$ from covariance $\langle \delta v_i(t)\delta v_j(t+\tau)\rangle = \mathbf{C}(\tau)$, we obtain

$$\frac{d}{d\tau}\mathbf{C}(\tau) = -\mathbf{C}(\mathbf{I} - \mathbf{W}^{\text{eff}})^T + \mathcal{A}(\tau)(\mathbf{W}^{\text{eff}})^T + \langle \delta v_i(t)\eta_j(t+\tau)\rangle \; . \tag{107}$$

This equation shows that the shared external noise generates the term $\langle \delta v_i(t)\eta_j(t+\tau)\rangle$ and thus provides the source of noise correlations in the model Eq. (102). In contrast, the network of binary units has no shared external noise, and the dynamics of correlations based on the master equation are:

$$\frac{d}{d\tau}\mathbf{C}(\tau) = -\mathbf{C}(\mathbf{I} - \mathbf{W}^{\text{eff}})^T + \mathcal{A}(\tau)(\mathbf{W}^{\text{eff}})^T \; . \tag{108}$$

As an approximation, we assume that the response of the rate variable $v_i$ to noise is quasi-static, so that $\langle \delta v_i(t)\eta_j(t+\tau)\rangle$ can be estimated as

$$\begin{aligned}\langle \delta v_i(t)\eta_j(t+\tau)\rangle &= \langle [\frac{d}{dt} + \mathbf{I} - \mathbf{W}^{\text{eff}}]^{-1}_{ik}\eta_k(t)\eta_j(t+\tau)\rangle \\ &= \int dw[iw + \mathbf{I} - \mathbf{W}^{\text{eff}}]^{-1}_{ik}\langle \tilde{\eta}_k(w)\, \tilde{\eta}_j(-w)\rangle \exp(iw\tau) \\ &\approx [\mathbf{I} - \mathbf{W}^{\text{eff}}]^{-1}_{ik}\langle \eta_k(t)\eta_j(t+\tau)\rangle \\ &\approx \left([\mathbf{I} - \mathbf{W}^{\text{eff}}]^{-1}\mathbf{\Sigma}_0\right)_{ij}\exp(-\frac{\tau}{\tau_0}) \; . \end{aligned} \tag{109}$$

We can also derive the equal-time correlation:

$$\langle \delta v_i(t)\eta_j(t)\rangle \approx \left([\mathbf{I} - \mathbf{W}^{\text{eff}}]^{-1}\mathbf{\Sigma}_0\right)_{ij} \; . \tag{110}$$

$$\langle \eta_i(t)\delta v_j(t)\rangle \approx \left(\mathbf{\Sigma}_0\left([\mathbf{I}-\mathbf{W}^{\text{eff}}]^{-1}\right)^{\text{T}}\right)_{ij} \,. \tag{111}$$

The time-evolution of equal-time correlation function $\langle \delta v_i(t)\delta v_j(t)\rangle$ is given by

$$\frac{d}{dt}\langle \delta v_i(t)\delta v_j(t)\rangle = -\langle \delta v_i(\mathbf{I}-\mathbf{W}^{\text{eff}})_{jk}\delta v_k\rangle - \langle \delta v_k(\mathbf{I}-\mathbf{W}^{\text{eff}})_{ik}\delta v_j\rangle + \langle \delta v_i\eta_j\rangle + \langle \eta_i\delta v_j\rangle \,. \tag{112}$$

In the matrix form, it can be written as

$$\frac{d}{dt}\mathbf{C}(t) = -\mathbf{C}(\mathbf{I}-\mathbf{W}^{\text{eff}})^T-(\mathbf{I}-\mathbf{W}^{\text{eff}})\mathbf{C}+\mathcal{A}(0)(\mathbf{W}^{\text{eff}})^T+\mathbf{W}^{\text{eff}}\mathcal{A}(0)+\langle \delta v_i\eta_j\rangle+\langle \eta_i\delta v_j\rangle \,, \tag{113}$$

where for convenience we separated the variance $\langle \delta v_i(0)\delta v_i(0)\rangle = \mathcal{A}(0)$ from the covariance $\mathbf{C}$. Substituting Eq. (110) and Eq. (111) into Eq. (113), we obtain

$$\begin{aligned}\frac{d}{dt}\mathbf{C}(t) \;&\approx\; -\mathbf{C}(\mathbf{I}-\mathbf{W}^{\text{eff}})^T - (\mathbf{I}-\mathbf{W}^{\text{eff}})\mathbf{C} + \mathcal{A}(0)(\mathbf{W}^{\text{eff}})^T + \mathbf{W}^{\text{eff}}\mathcal{A}(0)\\ &+\; [\mathbf{I}-\mathbf{W}^{\text{eff}}]^{-1}\mathbf{\Sigma}_0 + \mathbf{\Sigma}_0\left([\mathbf{I}-\mathbf{W}^{\text{eff}}]^{-1}\right)^{\text{T}} \,. \end{aligned} \tag{114}$$

The steady state solution of this equation is given by

$$\begin{aligned} 0 \;&=\; -\mathbf{C}(\mathbf{I}-\mathbf{W}^{\text{eff}})^T - (\mathbf{I}-\mathbf{W}^{\text{eff}})\mathbf{C} + \mathcal{A}(0)(\mathbf{W}^{\text{eff}})^T\\ &+\; \mathbf{W}^{\text{eff}}\mathcal{A}(0) + [\mathbf{I}-\mathbf{W}^{\text{eff}}]^{-1}\mathbf{\Sigma}_0 + \mathbf{\Sigma}_0\left([\mathbf{I}-\mathbf{W}^{\text{eff}}]^{-1}\right)^{\text{T}} \,. \end{aligned} \tag{115}$$

For the binary-unit model, we can use the master equation formalism to derive the correlation matrix [53], which is the same as Eq. (115) but without the input noise $\mathbf{\Sigma}_0$:

$$0 = -\mathbf{C}(\mathbf{I}-\mathbf{W}^{\text{eff}})^T - (\mathbf{I}-\mathbf{W}^{\text{eff}})\mathbf{C} + \mathcal{A}(0)(\mathbf{W}^{\text{eff}})^T + \mathbf{W}^{\text{eff}}\mathcal{A}(0) \,. \tag{116}$$

The equivalence of two forms of equations was derived in [54].

In the next subsections, we demonstrate that in the model with bistability and in the firing-rate model with a single fixed point and external noise [25], an attentional input, modeled as an input current $I$, changes the effective interaction strength leading to changes in correlations. In the network model with chaotic instability, an attentional input quenches the instability at specific spatial frequencies and thereby suppresses correlations by reducing the internal source of variability.

#### 3.2 Structure of neural correlations in the network with bistability

In the firing-rate model with bistability, the correlation of firing rates is proportional to the correlation of binary states:

$$r_i = \Delta r_i S_i + r_{\text{off},i} \,, \qquad \text{cov}(r_i, r_j) = \Delta r_i \Delta r_j C_{ij} \propto C_{ij} \,, \tag{117}$$

where $C_{ij} = \text{cov}(S_i, S_j)$. The steady-state solution for the correlation function is described by Eq. (116), which yields an approximate relation for the average $C$:

$$C \sim \frac{W^{\text{eff}}}{1 - (\sum)W^{\text{eff}}}\mathcal{A}(0) \; . \tag{118}$$

Here $(\sum)W^{\text{eff}}$ is the local interaction term, which only involves summation of nearby units, and therefore there is no scaling of $1/N$ in the interaction strength $W^{\text{eff}}$. Eq. (118) shows that $C$ is positively correlated with $W^{\text{eff}}$. In our model, $W^{\text{eff}} = \beta_1/(\alpha_1 + \alpha_2)$.

In the continuum limit, the interaction term becomes

$$(\sum)W^{\text{eff}} = \frac{\beta_1}{\alpha_1 + \alpha_2}(S_{x,y+1} + S_{x,y-1} + S_{x+1,y} + S_{x-1,y} - 4S_{x,y}) \to \frac{\beta_1}{\alpha_1 + \alpha_2}(\Delta d)^2 \cdot \nabla^2 \; , \tag{119}$$

In this case, the solution of Eq. (118) can be written as

$$C(d) = \exp\left(-\frac{d/\Delta d}{\xi}\right)\mathcal{A}(0) \; , \tag{120}$$

where the correlation length $\xi = \sqrt{\beta_1/(\alpha_1 + \alpha_2)}$. In this model, $\mathcal{A}(0)$ describes local bistable state transitions induced by independent noise in each unit, which is defined by intrinsic transition rates $\alpha_1$ and $\alpha_2$. The local metastability generates the source of variability, which propagates through the network via local spatial interactions.

In the network model with bistability, attentional input is modeled as a constant input current, which mainly changes the interaction strength $\beta_1$. The underlying mechanism for the attention-induced change in noise correlations is related to the non-linearity of the transition rate:

$$w(0 \to 1) = \frac{1}{\tau_{\text{off}}} = \frac{1}{A_1 + B_1 \exp[-C_1(2W \sum S_i + I)]} \approx \alpha_1 + \beta_1(\sum S_i) \; . \tag{121}$$

The interaction strength $\beta_1$ is the first-order derivative of the transition rate $w(0 \to 1)$ with linearized approximation. In the relevant parameter region, $w(0 \to 1)$ is a monotonically increasing and sub-linear function of the total input (which includes recurrent input $(\sum S_i)$ and external input $I$):

$$\beta_1 = \frac{\partial w(0 \to 1)}{\partial(\sum S_i)}|_I > 0 \; , \quad \frac{\partial \beta_1}{\partial I} \propto \frac{\partial \beta_1}{\partial(\sum S_i)} = \frac{\partial^2 w(0 \to 1)}{\partial^2(\sum S_i)} < 0 \; . \tag{122}$$

This fact implies that with increasing external input $I$, $w(0 \to 1)$ is closer to the saturation regime, where the dependence of $w(0 \to 1)$ on the input current $(\sum S_i)$ is weaker. In other words, the second derivative of $w(0 \to 1)$ with respect to $(\sum S_i)$ is less than zero, hence the interaction strength $\beta_1$ decreases with increasing external input current $I$. Therefore, the transition from Off to On states is less influenced by the recurrent inputs ($\sum S_i$), which underlies the reduction of correlations with increasing external input.

With increasing input current $\Delta I > 0$, the change in effective interaction strength $\Delta W^{\text{eff}}$ is approximately proportional to changes in $\beta_1$ (neglecting changes in $\alpha_1$ and $\alpha_2$, which are only moderate in the relevant parameter region):

$$\begin{aligned}\Delta W^{\text{eff}} &\approx \frac{\Delta\beta_1}{\alpha_1+\alpha_2} = \frac{1}{\alpha_1+\alpha_2}\left[\frac{\partial w(0\to 1)}{\partial(\sum S_i)}|_{I+\Delta I} - \frac{\partial w(0\to 1)}{\partial(\sum S_i)}|_{I}\right] \\ &\propto \frac{1}{\alpha_1+\alpha_2}\left[\frac{\partial^2 w(0\to 1)}{\partial^2(\sum S_i)}\right]\Delta I < 0\ , \quad (\text{if } \Delta I > 0)\ . \end{aligned} \tag{123}$$

Therefore, our model predicts the decrease of $W^{\text{eff}}$ with positive $\Delta I$. Since $C$ is an increasing function of $W^{\text{eff}}$, we expect a reduction of noise correlations:

$$\Delta C \sim \Delta\left[\left(\frac{W^{\text{eff}}}{1-(\sum)W^{\text{eff}}}\right)\mathcal{A}(0)\right] = \left(\Delta\frac{W^{\text{eff}}}{1-(\sum)W^{\text{eff}}}\right)\mathcal{A}(0)\ . \tag{124}$$

Notice that the autocorrelation $\mathcal{A}(0)$, which is the source of variability, does not change significantly, since $\mathcal{A}(0)$ only dependents on $\alpha_1$ and $\alpha_2$. The changes in correlation $\Delta C$ are mainly induced by $W^{\text{eff}}$. This mechanism is different from the model with chaotic instability [24], where suppression of variability $\mathcal{A}(0)$ at particular spatial frequencies underlies the reduction of correlations $\Delta C$ (see section 3.3).

In the Fourier space, the change of correlations can be written as

$$\Delta C(k) \propto \frac{1}{\Delta(1/W_{\text{eff}}) + (\Delta d)^2 k^2}\mathcal{A}(0) = \frac{1}{\Delta(1/\xi^2) + (\Delta d)^2 k^2}\mathcal{A}(0)\ . \tag{125}$$

Here we see that changes of the interaction strength $W^{\text{eff}}$ are equivalent to changes of the correlation length $\xi$, which affect different Fourier modes in a hierarchical manner. The amplitude changes decrease with increasing wave-number $k$ and vanish in the large $k$ limit. Summing these changes and performing the inverse Fourier transform, we find

$$\xi \to \xi - \Delta\xi\ , \quad \Delta C(d) = \left[\exp\left(-\frac{d}{\xi - \Delta\xi}\right) - \exp\left(-\frac{d}{\xi}\right)\right]\mathcal{A}(0)\ . \tag{126}$$

This result shows that the change in correlations is a non-monotonical function of distance $d$ that peaks at a finite distance.

#### 3.3 Structure of neural correlations in the network with chaotic instability

The spiking network model with internally generated variability [24] has no shared external input noise, so that $\mathbf{\Sigma}_0 = 0$ and the steady-state solution for the correlation function is described by Eq. (116). Here the variability of spiking is contained in the term $\mathcal{A}(0)$, which effectively provides the source of variability in corresponding firing-rate model. Considering one dimension in Eq. (116), which can be thought of as one Fourier mode, we find approximately

$$C \sim \frac{W^{\text{eff}}}{1-(\sum)W^{\text{eff}}}\mathcal{A}(0)\ . \tag{127}$$

For the spiking network with random spare connectivity [36, 53], $W_{\text{eff}} = \frac{1}{N}\tilde{W}_{\text{eff}}$, and $(\sum)W_{\text{eff}} = \tilde{W}_{\text{eff}}$, so that

$$C \sim \frac{\tilde{W}^{\text{eff}}}{1-\tilde{W}^{\text{eff}}}\frac{1}{N}\mathcal{A}(0)\ . \tag{128}$$

When the global activity reaches a fixed point, in the weak interaction limit or the E-I balanced regime, $\tilde{W}_{\text{eff}} \ll 1$, the correlation $C \sim \mathcal{A}(0)/N$ [36, 53], i.e. it vanishes at the large $N$ limit. On the other hand, in the spiking network with spatially local connectivity [24], at small wave-number Fourier modes, the mean activity stays in the unstable regime, which amplifies the variability $\mathcal{A}(0)$ and creates large correlations in these Fourier modes. The visual attention signal is modeled as a static depolarizing current to inhibitory neurons, which stabilizes activity at zero frequency Fourier mode. Therefore, for this mode

$$\Delta C \sim \frac{W^{\text{eff}}}{1-(\sum)W^{\text{eff}}}\Delta\mathcal{A}(0)\ , \tag{129}$$

where $\Delta\mathcal{A}(0)$ formally describes the reduction of variability from instability regime to a stable fixed point. So the reduction of noise correlation is mainly spatially homogeneous, corresponding to zero frequency Fourier mode.

#### 3.4 Structure of neural correlations in the network with externally driven fluctuation around a stable fixed point

The firing rate model with a single stable fixed point [25] has no internal source of variability $\mathcal{A}(0)$. To generate variability, the model assumes a correlated external input noise $\mathbf{\Sigma}_0$, so that the network operates just as a spatial filter of the external noise. The spatiotemporal pattern of input noise $\mathbf{\Sigma}_0$, shaped by the network filter, produces observed patterns of correlations. In this case, the dynamical equation is analogous to Langevin equation, and interaction terms acts as viscous force.

The steady-state correlation is given by Eq. (115) after absorbing $\mathcal{A}(0)$ into $\mathbf{C}$:

$$0 = -\mathbf{C}(\mathbf{I}-\mathbf{W}^{\text{eff}})^T - (\mathbf{I}-\mathbf{W}^{\text{eff}})\mathbf{C} + [\mathbf{I}-\mathbf{W}^{\text{eff}}]^{-1}\mathbf{\Sigma}_0 + \mathbf{\Sigma}_0\left([\mathbf{I}-\mathbf{W}^{\text{eff}}]^{-1}\right)^{\text{T}}\ . \tag{130}$$

The solution of this equation is

$$\mathbf{C} = (\mathbf{I}-\mathbf{W}^{\text{eff}})^{-1}\mathbf{\Sigma}_0\left([\mathbf{I}-\mathbf{W}^{\text{eff}}]^{-1}\right)^{\text{T}}\ . \tag{131}$$

The equation of the same form has been derived in different ways [55, 56, 57, 58]. This equation shows that the interaction term $(\mathbf{I}-\mathbf{W}^{\text{eff}})$ acts as a viscous force that suppresses the transformation of the input noise structure. Without interactions, the network becomes an ideal filter that inherits original patterns of shared input: $\mathbf{C} \to \mathbf{\Sigma}_0$. With the interaction term, the correlation involves a sum of inverse of $\lambda^2$, where $\lambda$ are eigenvalues of the matrix $(\mathbf{I} - \mathbf{W}^{\text{eff}})$ [25]:

$$\mathbf{C} \approx \left( \frac{1}{|\lambda|^2} + \dots \right) \mathbf{\Sigma}_0 \,, \tag{132}$$

In this model, a large attentional input current $I$ increase the effective interaction strength $W_{\text{eff}}$, which leads to larger absolute eigenvalues $|\lambda|$ in the E-I network, hence larger suppression of the input noise $\mathbf{\Sigma}_0$ [25]. Therefore, the correlation $\mathbf{C}$ is reduced with large attentional input current $I$. According to this analysis [25], when the external current is large, $|\lambda| \sim \sqrt{I}$, therefore $\mathbf{C} \sim (1/|\lambda|^2)\mathbf{\Sigma}_0 \propto 1/I$. The spatial structure of the changes in correlations depends on the combination of the spatial profile of $1/|\lambda|^2$ and the spatial pattern of shared input fluctuations $\mathbf{\Sigma}_0$:

$$\Delta\mathbf{C} \approx - \left( \frac{2\Delta|\lambda|}{|\lambda|^3} \right) \mathbf{\Sigma}_0 \,. \tag{133}$$

The mechanisms are fundamentally different between this model and the model with bistability. In the model with a single fixed point, an increasing input current increases the effective interaction strength and hence increases $|\lambda|$, which is analogous to increasing the viscous force that suppresses transmission of the external noise, Eq. (132). In contrast, in the model with bistability, an increasing input current reduces the effective interaction strength. According to Eq. (118), the effective interaction strength directly connects the internal local variability to correlations, so the reduction of effective interaction strength results in the decrease of correlations.

### 4 Supplementary figures

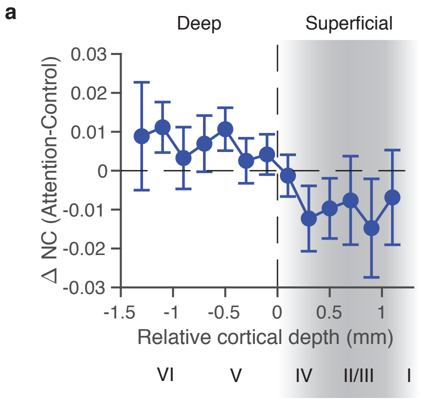

**Supplementary Figure 1. Changes in noise correlations during attention as a function of cortical depth.** Average change in noise correlations during attention as a function of the relative cortical depth. The relative cortical depth is defined to be zero at the boundary between the deep and superficial layers in the current source density measured in response to the full-field flash stimulus [26]. Probable locations of anatomical layers are shown below (Roman numerals). For each recording, we computed the average noise correlations for all possible ensembles of 6 nearby channels. For each ensemble, the average relative cortical depth was defined as the average of the 6 channels. We then grouped noise correlations based on the average relative cortical depth across all recordings.

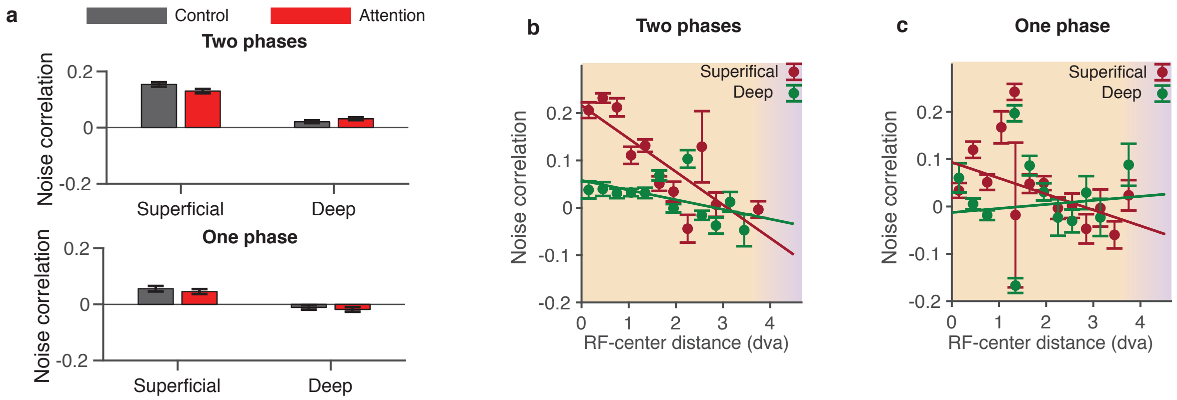

**Supplementary Figure 2. Dependence of noise correlations between single units (SUs) on the cortical layer, lateral distance and attention.** (**a**) Average noise correlation in superficial and deep layers in one-phase (upper panel) and two-phase (lower panel) recordings, separately for control (grey) and attention (red) conditions. (**b**) In two-phase recordings, noise correlations decrease with the RF-center distance in both superficial (crimson) and deep (green) layers (linear regression, one-sided t-test, slope $-0.07 \pm 0.02$, $p = 0.00007$ in superficial layers and slope $-0.02 \pm 0.01$, $p = 0.04$ in deep layers). Orange background highlights the range of short lateral distances within single or nearby columns. Purple background highlights longer lateral distances between distant columns, such as distances covered by a Utah array, which are outside the range of our laminar recordings. Error bars represent the standard error of the mean (SEM). (**c**) Same as b for one-phase recordings. In one-phase recordings, noise correlations do not consistently decrease with the RF-center distance (linear regression, one-sided t-test, slope $-0.03 \pm 0.01$, $p = 0.01$ in superficial layers and slope $0.01 \pm 0.02$, $p = 0.7$ in deep layers).

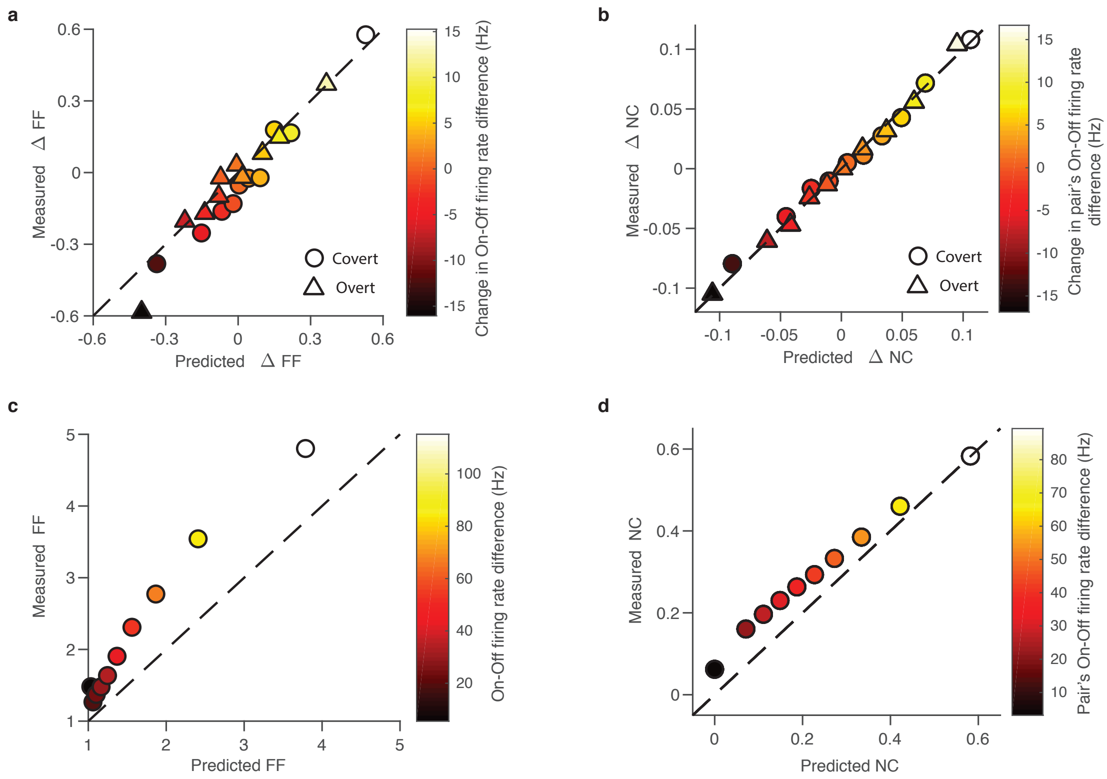

**Supplementary Figure 3. On-Off dynamics predict noise correlations within single columns.** (**a**) Comparison between covert(circle) and overt(triangle) attention-related changes in Fano factor ($\Delta\mathrm{FF} = \mathrm{FF}_{\mathrm{att}} - \mathrm{FF}_{\mathrm{ctl}}$) predicted by the On-Off model ($x$-axis) and measured from the data ($y$-axis). All neurons are divided in 10 equally-sized groups based on the change ($\Delta r_{\mathrm{att}} - \Delta r_{\mathrm{ctl}}$) in their On-Off firing-rate difference ($\Delta r = r_{\mathrm{on}} - r_{\mathrm{off}}$) between attention and control conditions. (**b**) Comparison between covert(circle) and overt (triangle) attention-related changes in noise correlations (($\Delta\mathrm{NC} = \mathrm{NC}_{\mathrm{att}} - \mathrm{NC}_{\mathrm{ctl}}$)) predicted by the On-Off model ($x$-axis) and measured from the data ($y$-axis). In this case, the pair's On-Off firing-rate difference is defined as $\sqrt{\Delta r_{\mathrm{att}_i} \Delta r_{\mathrm{att}_j}} - \sqrt{\Delta r_{\mathrm{ctl}_i} \Delta r_{\mathrm{ctl}_j}}$. (**c**) Comparison between Fano factor predicted by the On-Off model ($x$-axis) and measured from the data ($y$-axis). All neurons are divided in 10 equally-sized groups based on the On-Off firing rate difference. (**d**) Comparison between noise correlations (NC) predicted by the On-Off model ($x$-axis) and measured from the data ($y$-axis). All neurons are divided in 10 equally-sized groups based on the pair's On-Off firing rate difference.

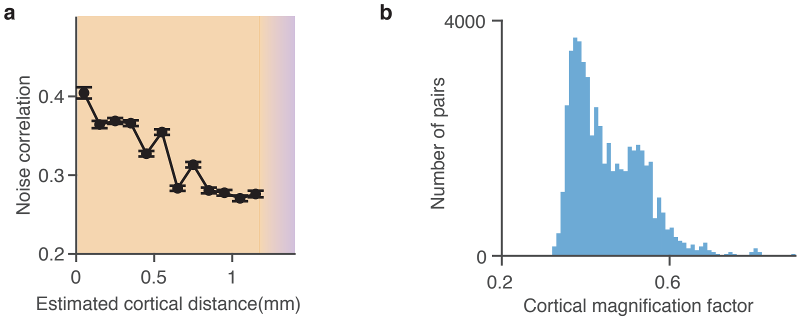

**Supplementary Figure 4. Average noise correlations decrease with the estimated cortical distance.** (**a**) Average noise correlations decrease with the estimated cortical distance, where cortical distance is estimated from the RF-center distance (dva) based on the cortical magnification factor. (**b**) Distribution of cortical magnification factors across all recording channels.
